## Supplementary Methods for "CNApp: quantification of genomic copy number alterations in cancer and integrative analysis to unravel clinical implications"

Symbol

^#^ Equal contribution.

^*^ Corresponding authors.

 (Camps J), (Castellví-Bel S).

**Supplementary methods**

Baseline adjustment of seg.mean values by sample purity

The amount of non-aberrant cell admixture may differ between cancer samples, necessitating separate adjustment of copy number thresholds for each assayed sample. To homogenize the analysis, purity estimations are used by CNApp to apply a baseline adjustment of the *seg.means* before the subsequent CNA calling by CNA amplitude thresholds. This adjustment allows for subsequent sample-to-sample comparison. *Seg.mean* values (*n*) by sample (*x*), when purity (*r*) available, are re-computed into new *seg.mean* values (*N*) as follows:

$$N(x)=2^{n(x)+(r(x)-1)}$$

When purity is not provided, 100% purity (r = 1) is assumed. Minimal purity accepted is 40% (r = 0.4), setting to 40% all purities below. Threshold of 40% also sets the negative capping value (C) as the maximal loss possible when heterozygous deletion happens (from 2 copies to 1 copy):

$$C = {log}_{2}\left( \frac{1}{2}\cdot0.4 \right)= -2.32$$

CNA scores computation (additional tables)

To compute CNA scores, copy number amplitude ranges are applied in order to weight focal and broad segments. Default amplitude thresholds to define those ranges are presented in the following table. CNA levels and absolute copy number are also specified:

| **Thresholds (Log2ratio values)** | **CNA level** | **Copy number** | **Weights (*A*)** |
| --- | --- | --- | --- |
| 1 | High-level gain | ≥ 4 copies | 3 |
| 0.58 | Medium-level gain | [3 – 4) copies | 2 |
| 0.2 | Low-level gain | [2.3 – 3] copies | 1 |
| -0.2 | Low-level loss | (1 – 1.7] copies | 1 |
| -1 | Medium-level loss | (0.6 – 1] copies | 2 |
| -1.74 | High-level loss | ≤ 0.6 copies | 3 |
| [-0.2 – 0.2] | Copy-neutral loss-of-heterozygosity (CN-LOH) | [1.7 – 2.3] copies  BAF ≥ 0.25 | 2 |

In order to take into account relative chromosome length from each segment when FCS is computed, coverage punctuation for focal events is implemented as specified in the following table:

| **%  chr-arm coverage** | **Coverage punctuation (*L*)** |
| --- | --- |
| ≤5% | 1 |
| >5% to ≤15% | 2 |
| >15% to ≤30% | 3 |
| >30% | 4 |

Correlation between CNA scores

Relation between Broad, Focal and Global CNA Scores (BCS, FCS and GCS, respectively) was assessed by computing Spearman’s rho statistic. The TCGA pan-cancer dataset including 10,635 samples spanning 33 cancer types was used to perform these associations. This data was downloaded from the GDC Data Transfer Tool from National Cancer Institute (Grossman et al., 2016).

Correlation between CNA scores and the fraction of genome altered

To evaluate the applicability of the CNA scores when it comes to sample CNA level assessment, we performed correlation analyses between CNA scores and the altered genome fraction. BCS and FCS were specifically correlated to the altered genome fraction by broad (i.e. chromosomal and arm-level events) and focal alterations, respectively.

We applied the re-segmentation procedure and CNA scores computation on a TCGA dataset comprised by 10,635 primary tumors from 33 TCGA projects analyzed by Affymetrix 6.0 SNP-array and DNACopy (Venkatraman & Olshen, 2007).

Global altered genome fraction (*altFract*) was computed, for each sample (*x*), by the summation of all copy number events lengths (*l*) and divided by total length of human genome (*hgLength*).

$$altFract(x)=\frac{\sum_{i=1}^{N} l_{i}}{hgLength}$$

Altered genome fractions for broad and focal events were computed, for each sample (*x*), by the summation of broad and focal alterations lengths (*l*), respectively, and divided by total length of human genome (*hgLength*).

$$Broad altFract (x)=\frac{\sum_{i=1}^{N} l_{i(Broad)}}{hgLength}$$

$$Focal altFract (x)=\frac{\sum_{i=1}^{N} l_{i(Focal)}}{hgLength}$$

Correlation test was performed by applying Spearman’s rho statistic to estimate a rank-based measure of association.

Associations between CNA scores and annotated variables

Annotated variables from input file are statistically associated with CNA scores computed by CNApp in *Re-Seg & Score* section. Different association tests are applied according to variable class (i.e. categoric or numerical), as described in Table 3. Both parametric and non-parametric tests are computed to present P-values in order to assess statistical significance for each association.

|  |  | *Parametric* | *Non-parametric* |
| --- | --- | --- | --- |
| **Categoric** | *n* = 2 | Student’s T-test | Wilcoxon signed-rank test |
|  | n > 2 | ANOVA | ANOVA: Kruskal test |
| **Numerical** |  | Pearson correlation | Spearman correlation |

*n* = groups defined by annotation variable

Survival analysis

Survival analysis by Kaplan-Meier was performed using *survival* and *survminer* R packages.

Correlation and clustering

Correlation and clustering methods can be applied into heatmap plots in CNApp *Region profile* section. Accepted correlation methods are: Pearson, Spearman and Kendall. Hierarchical cluster analysis is applied when clusters are computed in heatmaps.

**Supplementary Figure Legends**

**Figure S1 Correlation between CNA scores and the fraction of altered genome in the TCGA pan-cancer dataset**

Spearman’s rank correlation test was applied in order to assess the correlation between: **A**. Broad CNA Score (BCS) and fractions of altered genome affected by broad CNA events (whole chromosome and arm-level alterations) in each respective sample; **B**. Focal CNA Score (FCS) and the fraction of altered genome by focal CNA events in each respective sample; and **C**. Global CNA Score (GCS) and the overall fraction of altered genome in each respective sample. (TIFF 1.9MB)

**Figure S2 BCS and FCS dynamics TCGA pan-cancer dataset**

**A.** BCS values (x axis) and FCS values (y axis) scatterplot for 10.635 samples from pan-cancer cohort. Scatterplot smoothing by *loess* regression curve (in blue) and confidence interval; **B**. Correlation plot showing broad and focal CNA score values for the TCGA pan-cancer dataset. Spearman’s rank correlation test between broad and focal CNA scores (BCS and FCS, respectively) was performed. Sample subsets were computed by selecting 500 random samples from those displaying BCS ± 5 for each BCS value. Values of BCS and FCS for each subset of samples were correlated; **C**. Scatterplots for the 33 cancer types with BCS values (x axis) and FCS values (y axis) in samples. Scatterplot smoothing by *loess* regression curve (in blue) and confidence interval. (TIFF 3.1MB)

**Figure S3 Cancer types genomic profiles and clustering**

**A.** Heatmap analysis showing mean chromosome arm region profiles taking into account both broad and focal alterations, spanning 20 cancer type, for 8,653 samples from the TCGA pan-cancer dataset. Cancer types were indicated in the column *ID* from the input file. Annotation track displays tumor types, including squamous cell carcinomas, gynecological cancers, gastrointestinal cancers, and others; **B**. Heatmap analysis showing mean 5Mb region profiles taking into account only focal alterations, spanning 20 cancer type, for 8,653 samples from the TCGA pan-cancer dataset. Cancer types were indicated in the column *ID* from the input file. Annotation track displays tumor types, including squamous cell carcinomas, gynecological cancers, gastrointestinal cancers, and others; **C**. Heatmap plot showing 20 out of the 33 TCGA cancer type profile (by 5Mb genomic regions and accounting only for focal alterations) correlations, by Pearson's method, hierarchically clustered by tissue of origin. Gastrointestinal, gynecological and squamous cancers are clustering consistently in their respective groups. (TIFF 2.1MB)

**Figure S4 Frequencies of broad CNA events using different re-segmentation parameters**

**A.** The threshold of the relative length to the chromosome arm was set at ≥70% to classify segments as chromosome *arm-level*. **B**. Lowering the threshold of the relative length to the chromosome arm to ≥40%. **C**. Frequency plot using input data without the re-segmentation step. (TIFF 770KB)

**Figure S5 Analysis of CMS by FCS and differentially altered genomic regions**

**A**. CNA alterations frequencies of 462 colon cancer samples (TCGA-COAD cohort) using CNApp default thresholds for CNAs; **B**. Kaplan-Meier survival analysis across CMS groups for overall survival (see Supplementary Methods for clinical data extracted from TCGA data portal); **C**. Box plot showing the distribution of FCS according to the CMS groups for 309 colon adenocarcinoma samples. Wilcox test was performed for each pair of groups to assess significant differences. Significance is expressed as: p-value ≤ 0.001 (***); p -value ≤ 0.01 (**); p -value ≤ 0.05 (*); p -value > 0.05 (ns); **D**. BCS values density plot for MSI (brown) and MSS (green) sample groups after *Classifier model* section implementation selecting BCS as classifier variable (*N = 297* colon cancer samples). Vertical blue line at BCS = 4.75, as interception value between groups; **E.** Differentially altered chromosome arm regions between CMS groups. Heatmap plot displaying the significance between CMS groups paired comparisons. Student’s T-test was applied and multiple testing correction by BH method was used to assess differences in chromosome arm values between groups. Adjusted p-values are displayed. (TIFF 695KB)

**Supplementary Tables**

**Table S1:** Comparative analysis of broad CNA frequencies between CNApp and GISTIC2.0.(XLSX 15KB)

**Table S2:** Comparative analysis of focal CNA frequencies between CNApp and GISTIC2.0. (XLSX 70KB)

**Table S3:** Significant regional peaks identified by GISTIC2 in LIHC cohort. (XLSX 15KB)
