## Supplementary figures and images for "CNApp: quantification of genomic copy number alterations in cancer and integrative analysis to unravel clinical implications"

### Supplementary Figure S1

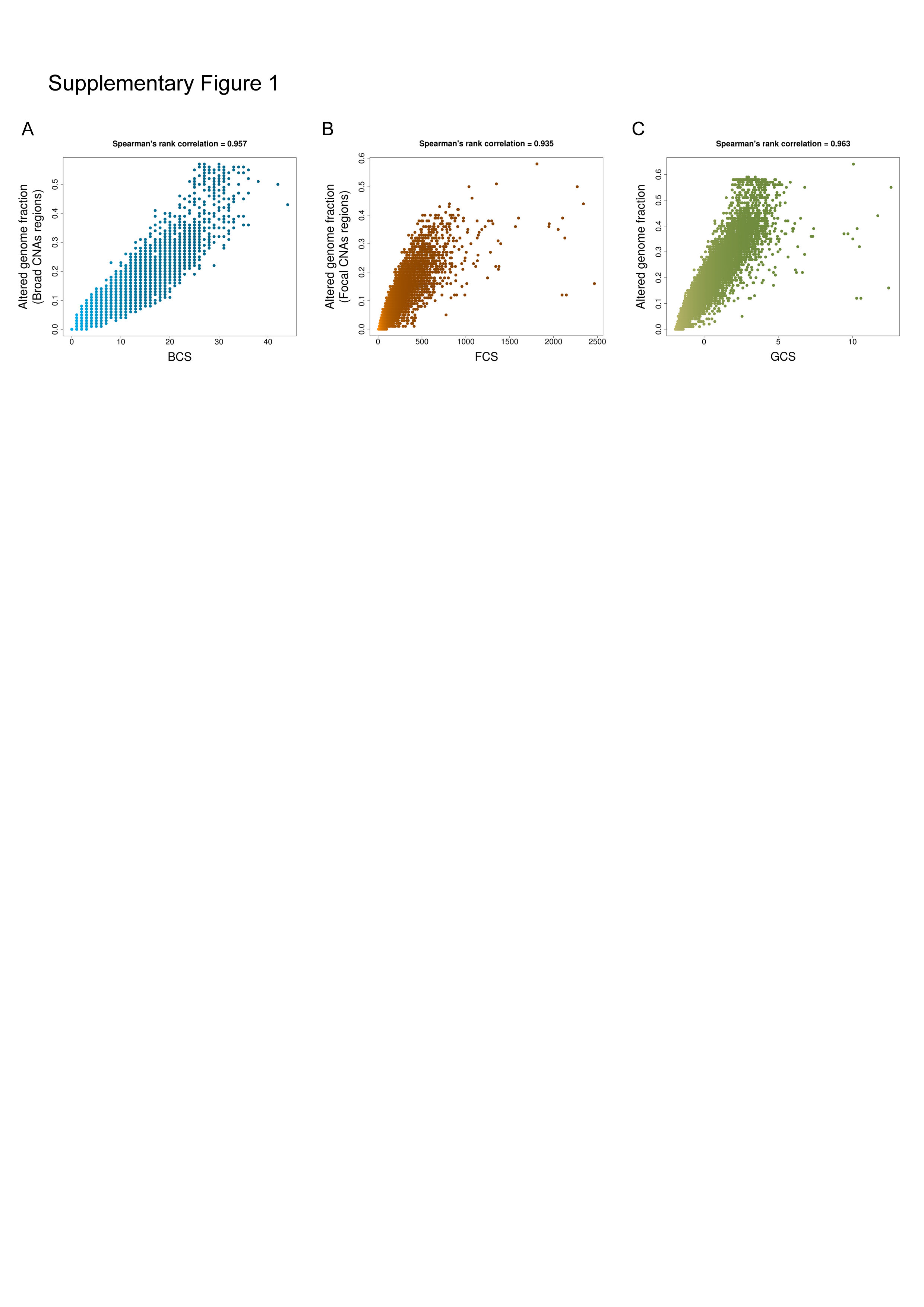

### Supplementary Figure S2

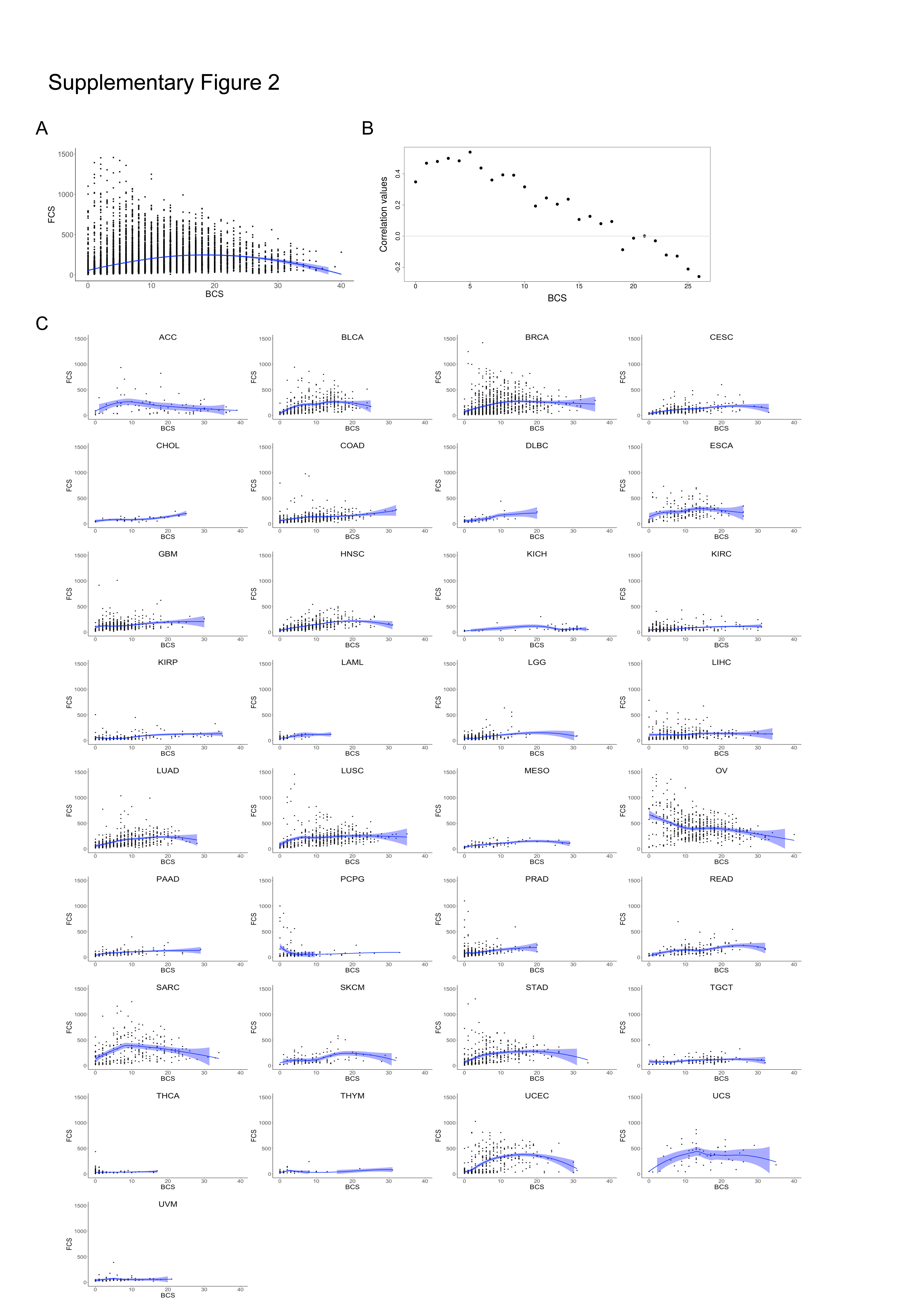

### Supplementary Figure S3

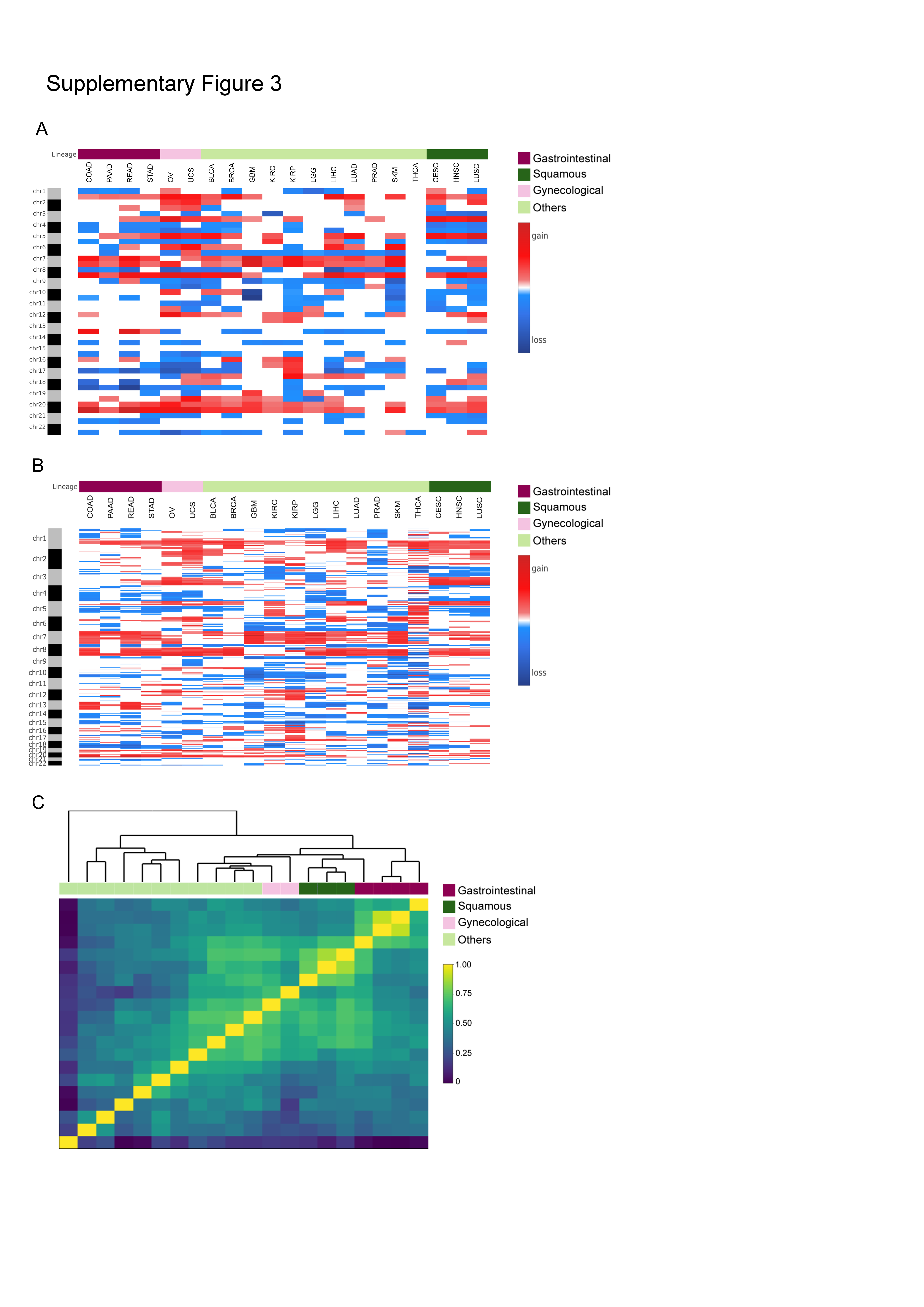

### Supplementary Figure S4

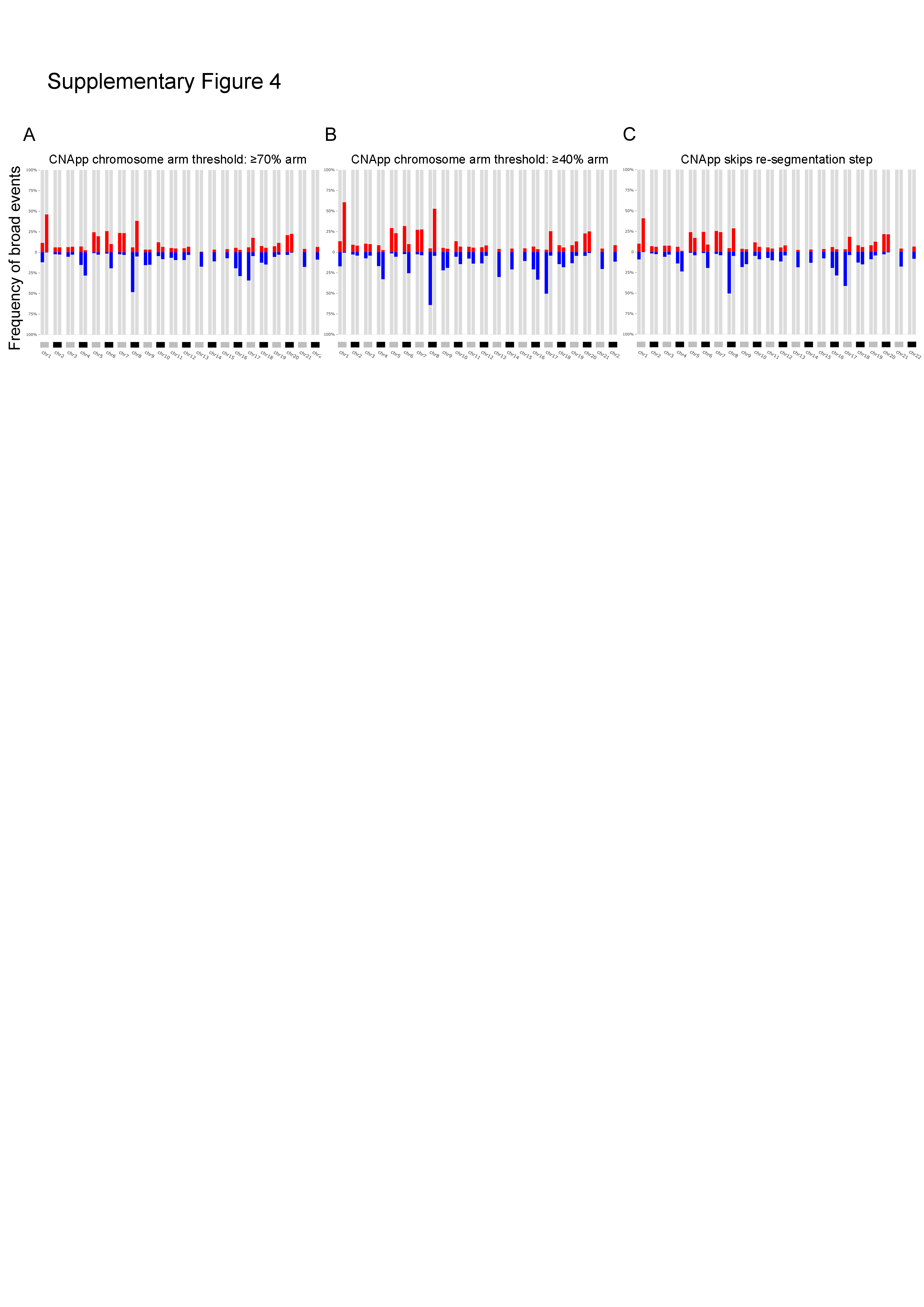

### Supplementary Figure S5

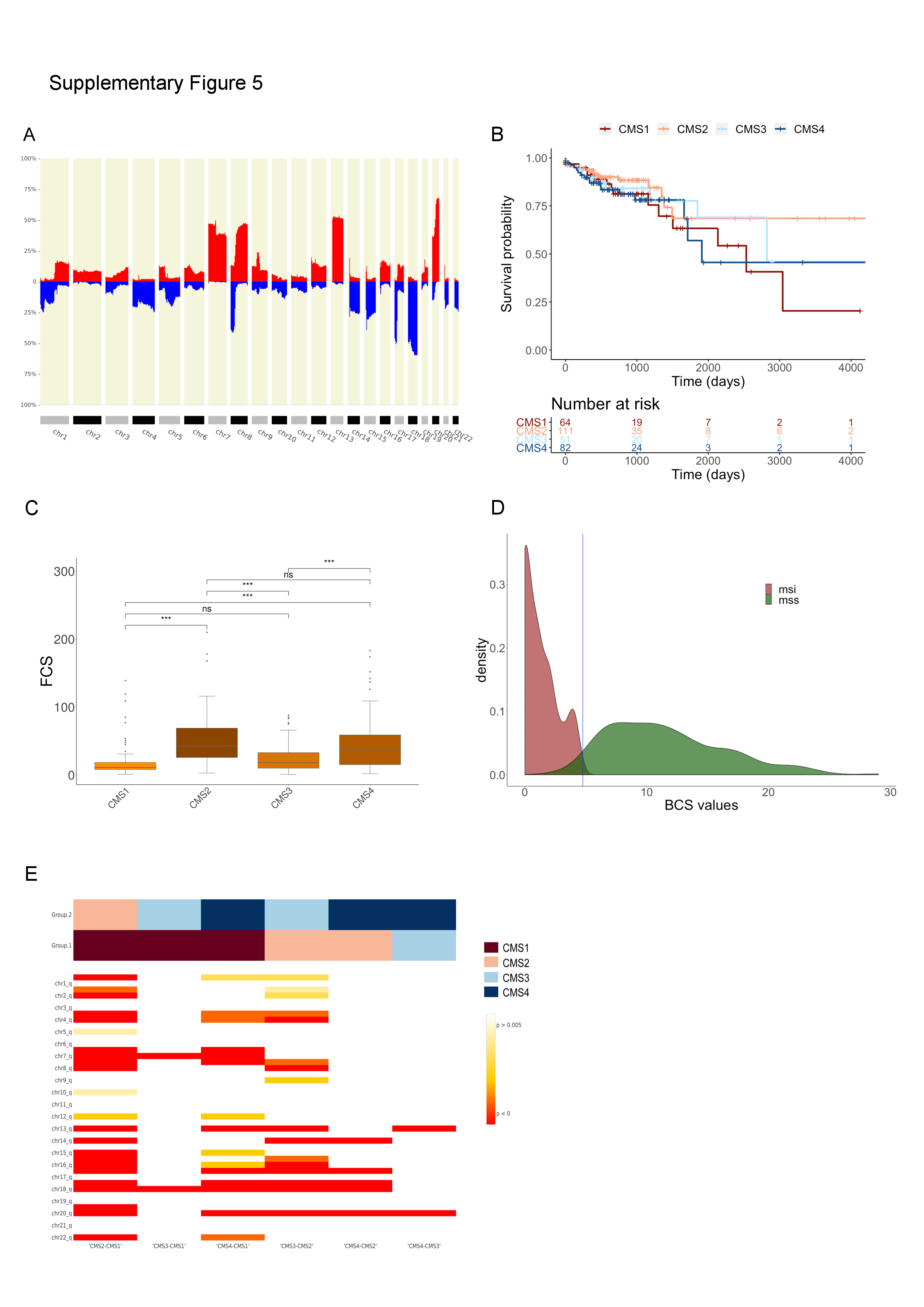
